# Respiratory Intervention Techniques Increase Selection Rate for Special Forces

**DOI:** 10.1101/774620

**Authors:** Bausek Nina, Summers Susanne, Scott B Sonnon

## Abstract

- Special Forces Selection Rates have declined over recent years, partly due to reduced fitness rates of applications
- Postural changes incurred prior to the recruitment may contribute to compromised respiratory function, resulting in fatigue, overexertion, and Selection Course failure
- Performance breathing training and respiratory muscle strengthening can reverse impaired respiratory function and optimize cardiopulmonary fitness.
- In this study, a 6-week intervention program including performance breathing training improved fitness and performance of SF Selection Course participants.
- The intervention program including performance breathing training increased the SF Selection Course pass rate from 0% to 30%, compared to prior years.

## Background

During intense sustained exercise, the inability to meet respiratory demand and exercise-induced dyspnea leads to physiological limitations and exercise cessation. Metabolic demands during high-intensity exercise are associated with extreme ventilation and gas exchange. In addition, the chemical stability of the blood is compromised during intense exercise, as ventilation is not able to buffer the blood pH anymore, contributing to carbon dioxide and lactic acid build-up. Furthermore, when the functional capacity of the respiratory system is reached, the respiratory muscle metaboreflex will compromise arterial oxygenation and limb oxygen flow [1].

Performance-defeating acquired breathing behaviors, poor respiratory muscle condition and compromised diaphragm function have a negative impact on physiology, psychology, health, and performance [2,3]. However, these behaviors can be deconditioned and reversed by specific countermeasures to improve performance, physiological health, and psychological status [4,5].

To qualify for selection into the Special Forces Unit, warriors are required to excel in physiologically and psychologically challenging environments. The number of candidates qualified for the Special Forces Unit selection process has declined in recent years, significantly decreasing the number of qualified candidates to fill the need our Armed Forces have [6].

The Special Forces Assessment and Selection (SFAS) course failure rate was four-times (4x) in 2017, compared to 2014 (passing rate in 2014: 12%, in 2017: 3%). In addition, SFAS candidates in the Army Physical Fitness Test (APFT) have five-times (5x) the failure rates, and a 22% increased injury rate, since 2014. (FY2014: Medical: 6.4%; APFT: 2.9% FY2017: Medical: 8.2%; APFT: 13.9%).

## Hypothesis

Warriors applying for SF recruitment often present with postural changes due to a variety of causes, including trauma, fixed positions, un-ergonomic gear carry, repetitive motions, suboptimal performance form/mechanics, sleep disruption, medication, and dietary changes. These can compromise the respiratory system and respiratory function, leading to faster rates of fatigue, increased overexertion injuries and illnesses, and increased Selection Course failure rates.

Changing these respiratory behaviors to reverse over-breathing patterns should result in increased physical performance, decreased injuries and as a result, increased selection course passing rates.

We investigated whether a 6-week intervention study during the SFAS course including RMT as well as respiratory and behavioral countermeasures improves physical performance and selection pass rate.

## Study Aim

The mission of the here described 6-week study is to help soldiers improve breathing strength and performance through the application of behavioral learning principles to breathing physiology during physical training in preparation for the *Special Forces Assessment and Selection Course*.

## Study Design

All subjects that enrolled in the study underwent a baseline assessment including pulmonary function testing (spirometry), respiratory muscle strength (maximum inspiratory pressure (MIP), maximum expiratory pressure (MEP), and exhaled carbon dioxide testing (EtCO2). All subjects also performed a 2-mile run.

For all subjects of group A, their capnic status at the anaerobic threshold was assessed during a loaded squat test.

Group A followed a 6-week intervention including two (2) sessions of RMT per day. Each RMT session consisted of two (2) sets of 10 breaths at an intensity setting corresponding to 70% of maximum effort. In order to avoid overtraining, subjects were instructed to reduce or skip sessions on days of strenuous exercise.

Furthermore, group A was instructed in a set of 3 types of Performance Breathing maneuvers and biofeedback through capnometry and pneumotagraphy (using CapnoTrainer and CapnoPlus):

1. Improve ventilation at the point of exercise cessation
2. To increase mindfulness
3. To improve the quantity and quality of sleep.

These fatigue countermeasures aimed to equip subjects in group A to withstand the physiological and psychological challenges presented during the SFAS course and the subsequent selection.

Subjects in group B completed the SFAS course alongside subjects from group A. Due to warfighting requirements, the endline assessment was limited to a 2-mile run and to a comparison of the selection pass rate to historical pass rate data.

## Results and Discussion

In total, 19 soldiers were enrolled in the study and were allocated to either group A (intervention group) or group B (control group).

Table 1 shows the demographics and baseline measurements of subjects enrolled in the study. The difference in demographics and baseline measurements between the two groups was assessed by ANOVA (StatPlus) and revealed no significant difference between the groups at baseline, with the exception of respiration rate, which showed a significant difference between group A and group B (highlighted by *). The observed difference here may be linked to activities prior to arrival at the assessment station, to stress or anxiety about the measurement methods, to lower levels of aerobic fitness, or to other reasons. As the differences in respiration rate were not reflected by other baseline parameters, they were disregarded for outcome analysis.

**Table 1:**
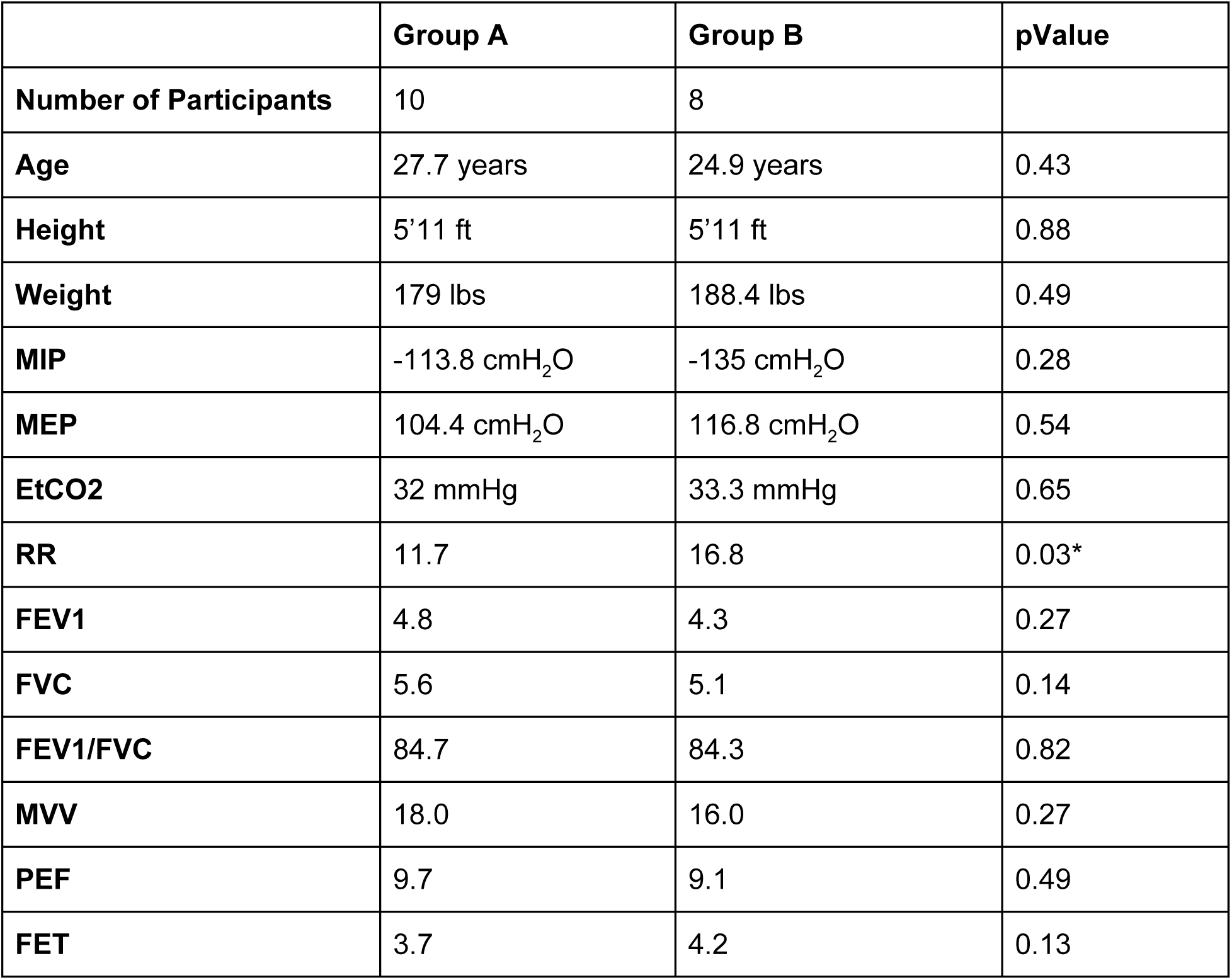
Soldier Demographics.

## Outcome Performance and Selection

All subjects performed a 2 mile run at baseline and endline of the study under comparable conditions and without load. Following the 6-week SFAS course, subjects from both groups underwent SF selection.

Tables 2 demonstrates the 2-mile run times after the study duration (Endline) were significantly different between the groups (highlighted by *, ANOVA, StatPlus). The average improvement in the intervention group A was 3.55%, while subjects in group B only improved by 0.52% on average. This difference in physical performance at the time of entering the special unit selection may have given warriors in group A a critical advantage, which is reflected in the selection rate. The selection rate in group A was at 30%, while none of the subjects in group B were selected. Figure 1, panel A depicts the significant difference in the average 2-mile run time between the groups (p=0.02, error bars represent standard deviation), while panel B shows the notable difference in average improvement in both groups. However, this difference in the percentage of improvement fails to reach significance, most likely due to the range of variability within the group (standard deviation in group A: 7.59, in group B: 9.62).

**Table 2:** Comparison of Major Outcomes at Baseline and End Line and Selection Rate.

|  | Baseline | End Line | % Improvement | % Selected |
| --- | --- | --- | --- | --- |
| **Intervention Group A** | 13:40 | 13:01 | 3.55% | 30% |
| **Control Group B** | 14:40 | 14:36 | 0.34% | 0% |
| **pValue** | 0.08 | 0.02* | 0.52 | N/A |

**Figure 1:**
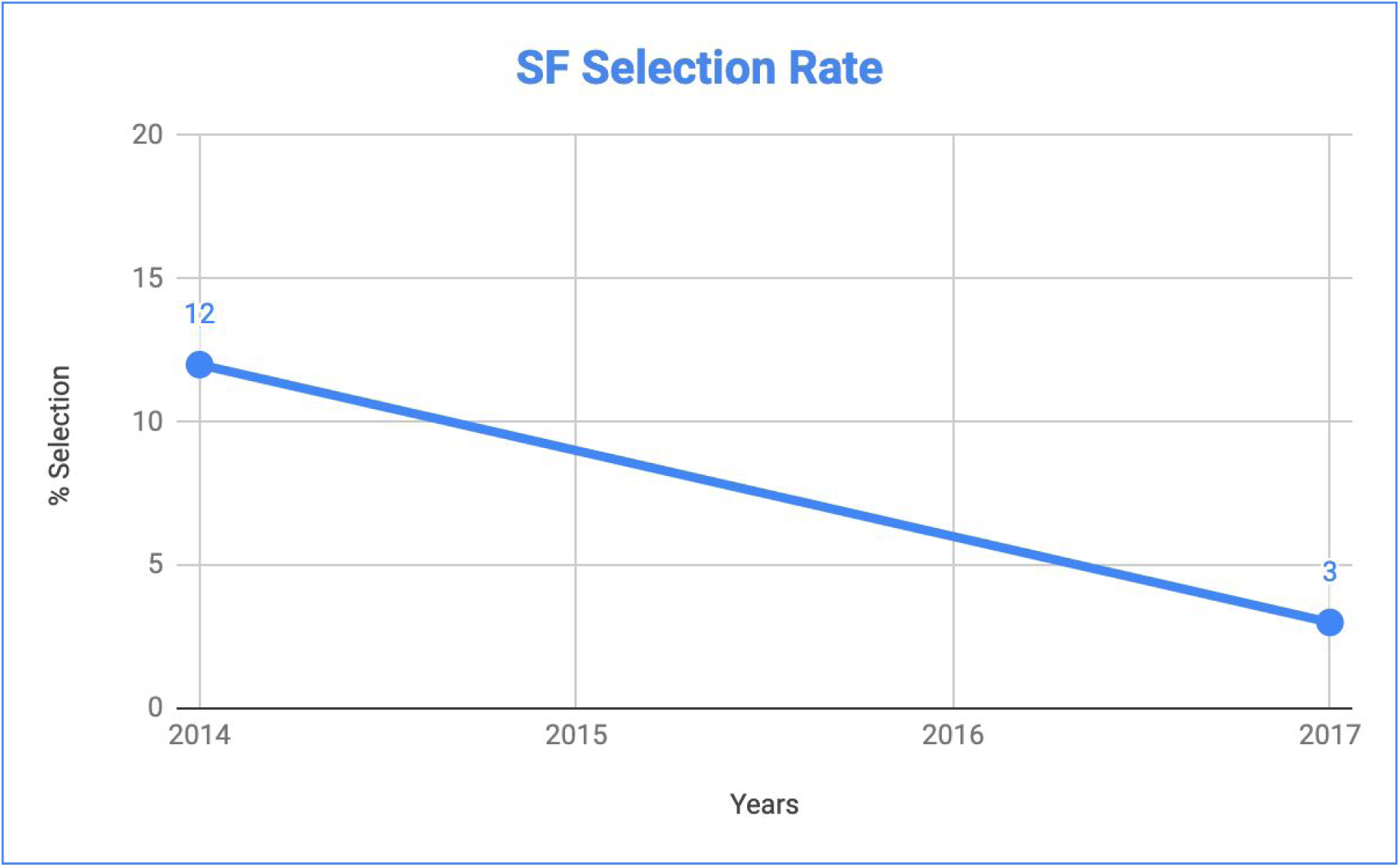
Historic Special Forces Selection Rate.

**Figure 1:**
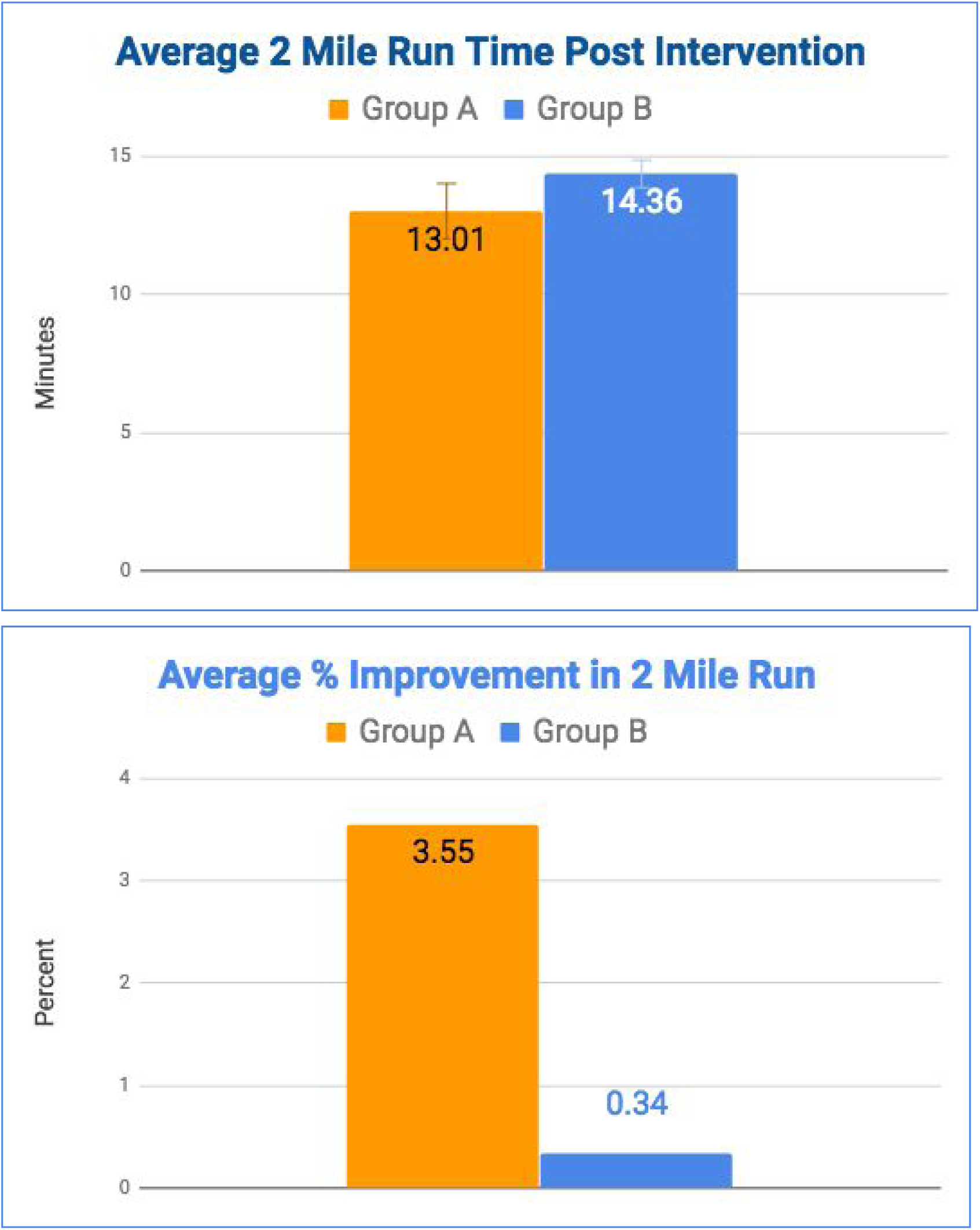
Average Endline (post intervention) Run Times and Improvement.

**Figure 2:**
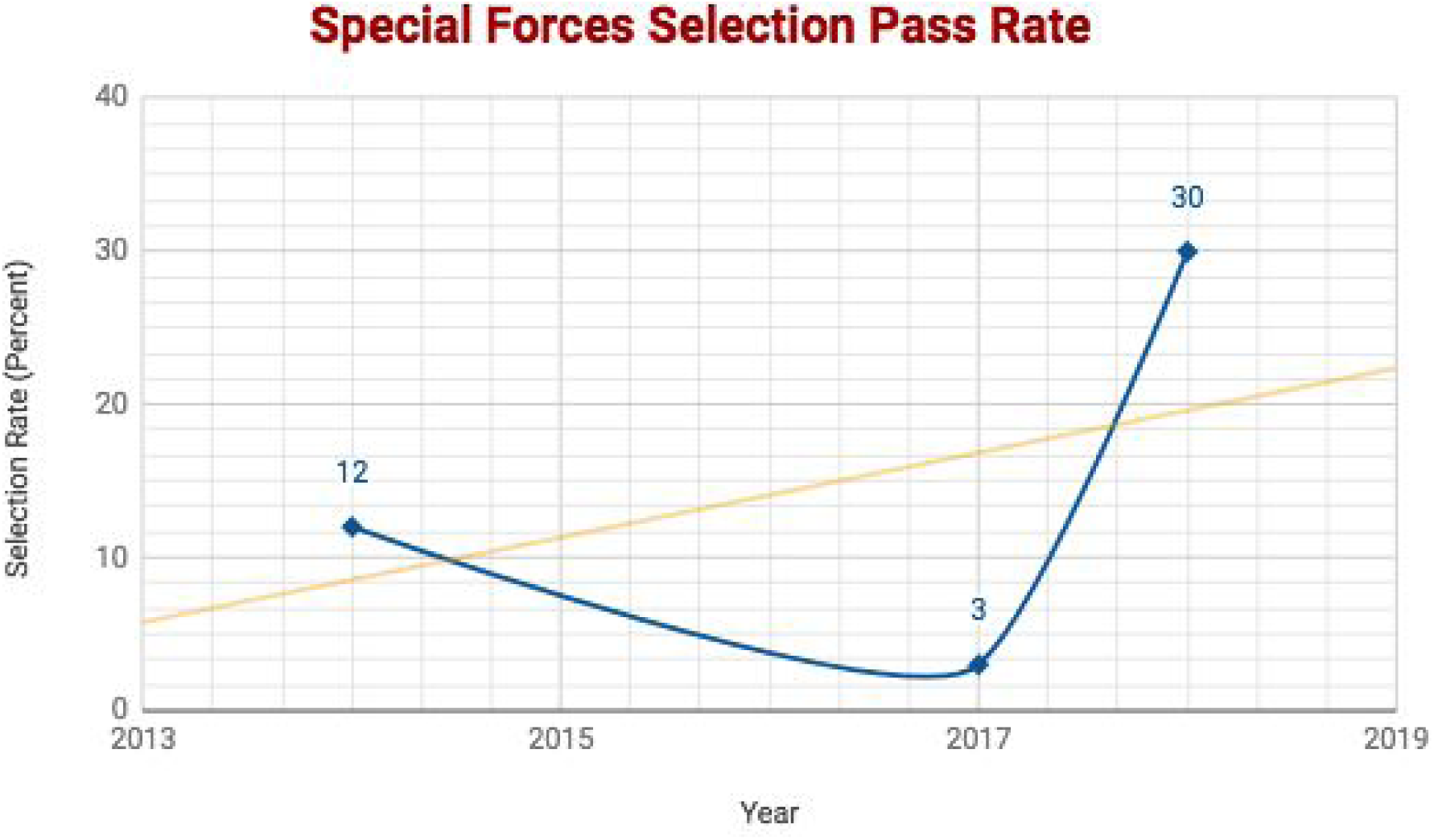
Selection Rate Compared to Historic Data and Trend.

## Conclusion

A 6-week intervention program including strengthening and preventative measures as well as fatigue countermeasures to improve physiological and psychological performance, resilience, and mindfulness was incorporated into the SFAS course to prepare for selection into special units. Resistive performance breathing training to strengthen the diaphragm and other respiratory muscles, as well as education in a situation-specific adjustment of breathing patterns were introduced to optimize respiratory function and to optimally prepare the cardiopulmonary system for the extreme demands during the selection process.

The intervention program was compared to the accomplishment of the SFAS course without adjunct intervention. We demonstrate that the intervention program including performance breathing training increases the average run time in the 2-mile time trial. We also show that the breathing training increases the selection rate to 30%, compared to 0% in the control group. Compared to historic selection rates from 2014 and 2017, the achieved selection rate of 30% could reverse a downward trend and increase selection rates for the future (orange trend line). However, the findings from this pilot study require confirmation in larger studies.

We, therefore, recommend the intervention program as an effective measure to support recruitment to Special Forces, and to enhance warrior resilience and performance.

### Disclosure

NB receives funding from PN Medical, makers of The Breather, due to work as their independent Chief Scientist. SS worked as a sales clinician for PN Medical.

## References

1. McConnell A. Respiratory Muscle Training: Theory and Practice. 1 edition. Churchill Livingstone; 2013.

2. Welch JF, Archiza B, Guenette JA, West CR, Sheel AW. Effect of diaphragm fatigue on subsequent exercise tolerance in healthy men and women. J Appl Physiol. 2018; doi: 10.1152/japplphysiol.00630.2018

3. Dominelli PB, Katayama K, Vermeulen TD, Stuckless TJR, Brown CV, Foster GE, et al. Work of breathing influences muscle sympathetic nerve activity during semi-recumbent cycle exercise. Acta Physiol. 2018; e13212.

4. Shei R-J. Recent Advancements in Our Understanding of the Ergogenic Effect of Respiratory Muscle Training in Healthy Humans: A Systematic Review. J Strength Cond Res. 2018;32: 2665–2676.

5. Downey AE, Chenoweth LM, Townsend DK, Ranum JD, Ferguson CS, Harms CA. Effects of inspiratory muscle training on exercise responses in normoxia and hypoxia. Respir Physiol Neurobiol. 2007;156: 137–146.

6. Balestrieri S. “I Went Thru the Last Hard Class” and The Ever Changing Standards I SpecialOperations.com “I Went Thru the Last Hard Class” and The Ever Changing Standards. In: SpecialOperations.com [Internet]. 24 Sep 2018 [cited 9 Sep 2019]. Available: https://specialoperations.com/33603/i-went-thru-the-last-hard-class-and-the-ever-changing-standards/

